# Deep Heterogeneous Dilation of LSTM for Transient-phase Gesture Prediction through High-density Electromyography: Towards Application in Neurorobotics

**DOI:** 10.1101/2021.10.26.466039

**Authors:** Tianyun Sun, Qin Hu, Jacqueline Libby, S. Farokh Atashzar

**Author notes:** Sun and Hu share the first authorship.

## Abstract

Deep networks have been recently proposed to estimate motor intention using conventional bipolar surface electromyography (sEMG) signals for myoelectric control of neurorobots. In this regard, Deepnets are generally challenged by long training times (affecting practicality and calibration), complex model architectures (affecting the predictability of the outcomes), and a large number of trainable parameters (increasing the need for big data). Capitalizing on our recent work on homogeneous temporal dilation in a Recurrent Neural Network (RNN) model, this paper proposes, for the first time, *heterogeneous temporal dilation* in an LSTM model and applies that to high-density surface electromyography (HD-sEMG), allowing for the decoding of dynamic temporal dependencies with tunable temporal foci. In this paper, a 128-channel HD-sEMG signal space is considered due to the potential for enhancing the spatiotemporal resolution of human-robot interfaces. Accordingly, this paper addresses a challenging motor intention decoding problem of neurorobots, namely, *transient intention identification*. Our approach uses only the dynamic and transient phase of gesture movements when the signals are not stabilized or plateaued, which can significantly enhance the temporal resolution of human-robot interfaces. This would eventually enhance seamless real-time implementations. Additionally, this paper introduces the concept of “dilation foci” to modulate the modeling of temporal variation in transient phases. In this work a high number (e.g., 65) of gestures is included, which adds to the complexity and significance of the understudied problem. Our results show state-of-the-art performance for gesture prediction in terms of accuracy, training time, and model convergence.

## I. Introduction

As of 2005, more than 1.6 million people in the United States were living with the loss of a biological limb. This population is estimated to double by 2050. Besides, accidents and congenital conditions, some medical conditions can lead to amputation, such as cancer, vascular diseases, diabetes, and peripheral arterial diseases [1]. The population of people who have such conditions is also growing in an accelerated manner. Thus, the research in fabrication and seamless control of prostheses is in substantially high demand. For upper-limb functions, due to the complexity and diversity of tasks, intuitive and agile (fast in response) control are technically challenging. Addressing these problems can help amputees with Activities of Daily Living (ADLs) beyond essential hand functions. Furthermore, existing gesture detection algorithms have low accuracy and high latency, leading to a high rejection rate in commercial systems [2]–[4].

Surface electromyography has been used extensively in the literature to implement myoelectric control of bionic limbs, allowing for peripheral interfacing of the human motor intention to robotic actions in a noninvasive manner [5]. sEMG-based gesture classification can be used as a reference for real-time robotic control. A conventional approach is to feed extracted temporal and spectral features from sEMG signals to classic models such as Support Vector Machines (SVMs) or Linear Discriminant Analysis (LDA) [6]–[8]. Researchers have also achieved high performance when feeding denoised sEMG from only two pairs of electrodes to a probabilistic classifier [9].

Deep learning techniques have been increasingly used to decode the complex human neurophysiological responses to motor commands, exploiting the rich information present in the sEMG signals. Convolutional neural networks (CNNs) have been leveraged in sEMG-based prosthetic studies [10]–[16] because of their ability to detect and localize human neurophysiological features in a given segment of muscle-activity signal. Recurrent Neural Networks have also been used [17]–[21] because they capture the underlying temporal dynamics from sEMG signals. Long-Short-Term Memory (LSTM) is a type of RNN [22]. An LSTM unit has a cell, an input gate, an output gate, and a forget gate, which work together to allow the model to capture both long and short dependencies of the signals.

Some recent articles [23],[24], including our previous work [25],[26], have proposed hybrid models that leverage the benefits of both CNNs and RNNs in gesture detection. In [25], we proposed a hybrid approach that achieves high performance on conventional user-specific and generalized gesture classification, with reduced need for re-training and re-calibration. However, traditional bipolar sEMG signals have challenges in capturing muscle group activities due to limited numbers of sensors and sparse sensor placement, thereby limiting the number of detected gestures. Most existing literature only uses the plateau phase of contraction, which is a steady-state phase during highly-controlled and instructed task conduction when the signal does not represent a dynamic contraction. The use of the steady segment of the signal results in low temporal resolution, late reaction, and incorrect classification during transient phases which can affect practicality and intuitiveness. This paper aims to address the aforementioned issues by proposing a new computational model that can process high-density surface electromyography (HD-sEMG) signals to enhance the spatiotemporal resolution of intention decoding.

High-density surface electromyography has attracted considerable attention in recent years because it encodes distributed activities of motor units across the muscles and the gradient of changes in time and space, which are critical factors for distinguishing intended motor tasks. HD-sEMG signals are noninvasively collected from a large number of electrodes arranged in a two-dimensional array. The dense placement (e.g., 5-10mm inter-electrode space) of electrodes in a 2D grid describes the muscle activities both as a function of time and topologically (in space) for the muscle group. Some recent efforts have been conducted to utilize various representations of HD-sEMG signals for detecting human intention. Examples are as follows: time-domain representation [27]–[29], image-based muscle activity heatmap representation [24],[27],[30], and motor unit action potentials and the corresponding spike trains derived through decomposition of HD-sEMG [31]–[33]. In the above literature, HD-sEMG has shown the ability to secure high accuracy. However, there are some critical limitations as follows: (a) signals are often down-sampled to reduce the volume of high-density information in some (not all) cases, (b) relatively low number of classified gestures (< 27 gestures) are considered, (c) low number of subjects, and (d) the plateaued phases of contraction is considered under controlled environments and long signal windows. In this paper, we use a new open dataset (see Section II-A), and specifically address the transient-phase decoding problem for a high number of gestures using the proposed novel algorithm. We conduct a comprehensive comparative study to support state-of-the-art results.

Despite the diversity of model structures (CNNs, RNNs, or hybrid models), the literature suffers from the most common deep-learning problems, including long training times, vanishing/exploding gradients, and short dependencies. Furthermore, these models suffer from traditional deep learning limitations, such as requiring large training sets to classify a large number of classes and avoid overfitting. Therefore, we previously proposed homogeneous temporal dilation by adding dilation into the LSTM module [26], modeling longer temporal dependencies and thereby mitigating vanishing/exploding gradients. At the same time, it makes the structure less complex, allowing the training time to be 20 times faster than existing counterparts.

In this paper, we are taking the next fundamental step by proposing nonlinear heterogeneous temporal dilation of a pure LSTM network to further explore the benefits of various dilation modes. In heterogeneous temporal dilation, the skipped LSTM cells exponentially increase within each layer, broadening the receptive field of the model for capturing longer and more diverse temporal dependencies. Additionally, we analyze the impact of dilation focus (see Fig. 4), which varies the connection density of the LSTM cells along the temporal dimension. Furthermore, we train and test on the transient phase of each repetition, which contains only 10% of the signal length. We also investigate the effect of different window lengths and achieve the best model performance with a window size of 200ms. The six contributions of this paper are summarized below:

**Fig. 1:**
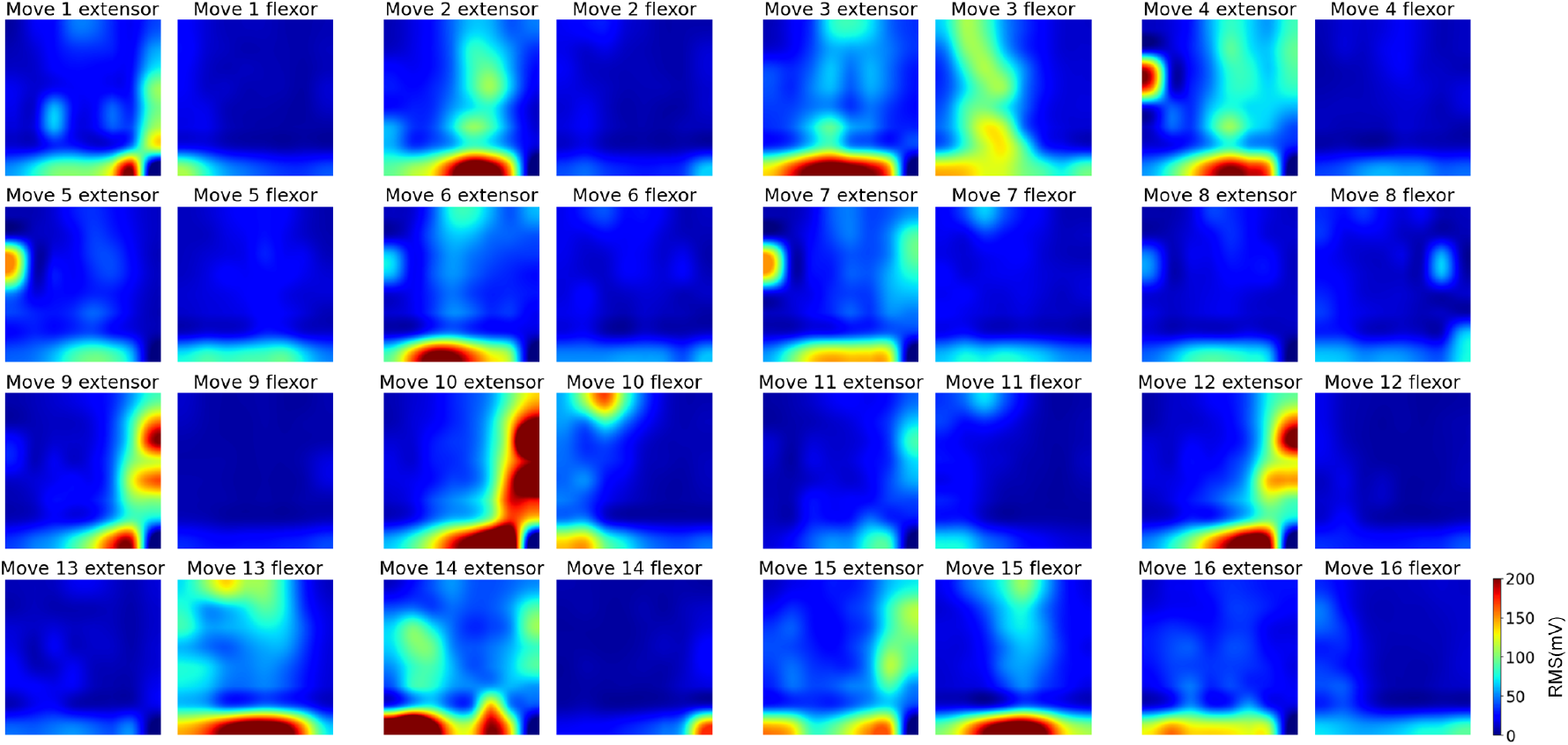
32 muscle-activity heatmaps associated with 16 1-DoF movements from the best-performing subject (#15). Each gesture has two heatmaps (forearm extensor and flexor). Each heatmap is an 8×8 grid, consisting 64 electrodes.

**Fig. 2:**
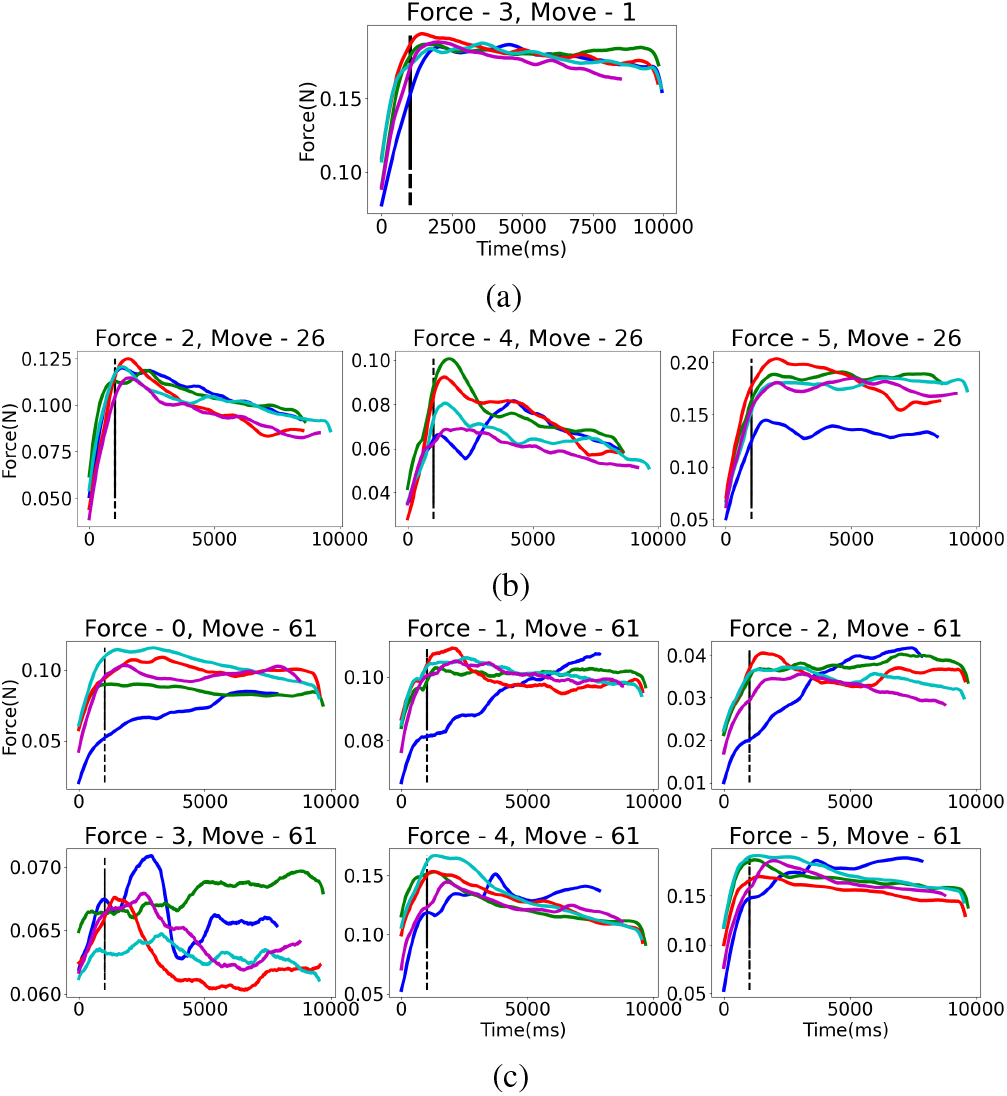
This figure shows the corresponding forces of three gestures with different DoFs on each repetition. The dashed lines indicate the end (0.5 seconds) of transient phases. Force indices 0-5 denote strain gauges on index finger, middle finger, ring finger, little finger, thumb finger flexion/extension, thumb finger abduction/adduction, respectively. Line colors denote five different repetitions. (a) Little finger force of little finger bend gesture; (b) Ring finger force and thumb forces of ring finger bend and thumb down gesture; (c) All five fingers forces of palmar grasp gesture.

**Fig. 3:**
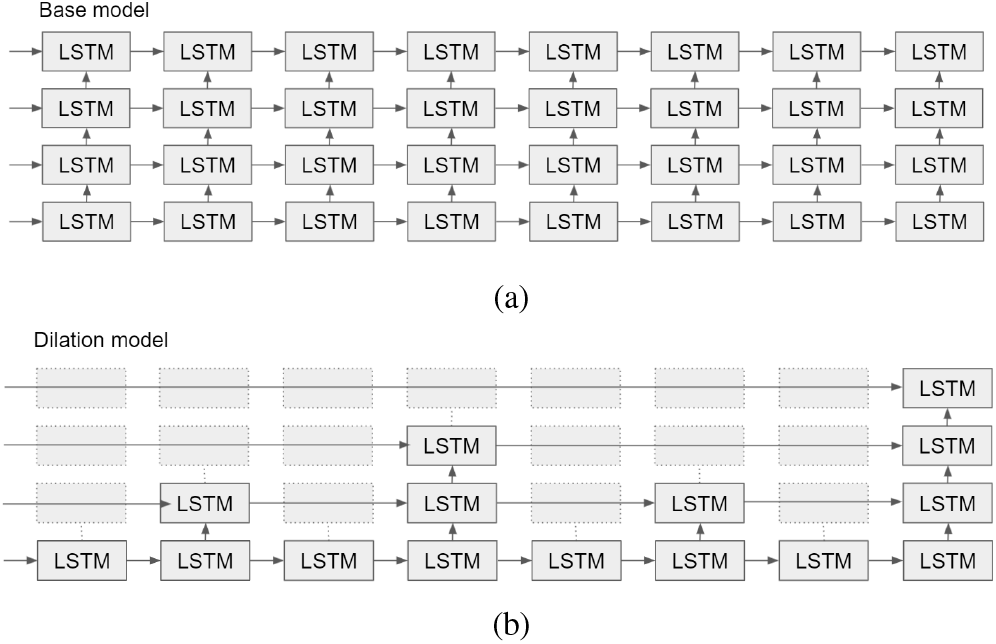
(a) Regular baseline model (all LSTM cells are connected); (b) Dilated baseline model with first-order dilation.

**Fig. 4:**
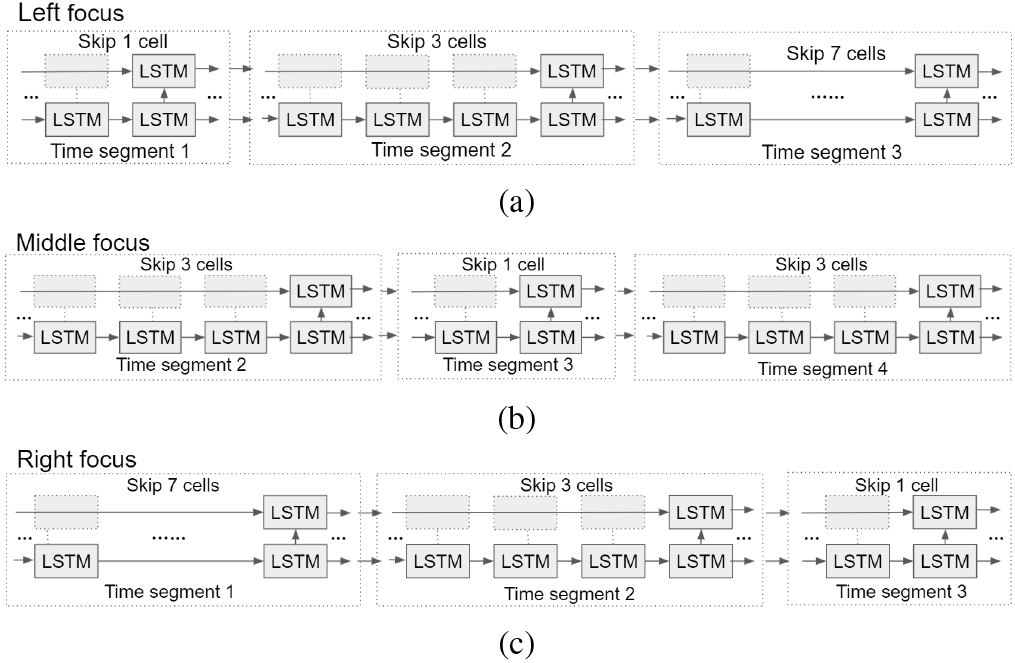
(a) Left-focused model where the highest connection density is at the beginning timestamps; (b) middle-focused model where the highest connection density is at the middle timestamps; (c) right-focused model where the highest connection density is at the end timestamps.

### Contribution 1

This paper proposes heterogeneous temporal dilation, for the first time, which introduces nonlinearity in the skip connections of LSTM cells to increase the reach and variety of temporal dependencies. The nonlinear dilation further alleviates deep learning problems such as vanishing/exploding gradients and long training times.

### Contribution 2

The proposed dilated model powerfully and successfully predicts a high number (65) of gestures, achieving 83% accuracy and enhancing the model versatility.

### Contribution 3

The gesture prediction task is designed to need only the transient phases of the signal (10% of the signal length from each repetition) which significantly enhances the agility and temporal resolution.

### Contribution 4

The concept of dilation foci is proposed and implemented for the first time, adding one more degree of freedom to the proposed Deepnet model, modulating the model’s temporal reach, which is beneficial for adaptability.

### Contribution 5

The analysis of varying window sizes found the best model performance (82% median accuracy) when the window size is 200ms (which is lower than the real-time requirement of 300ms in prosthesis control [34]).

### Contribution 6

The proposed heterogeneous dilation shortens the training time by more than 20 times over regular RNN models.

## II. Material and Methods

### A. Data Acquisition Process

In order to design a robust, lightweight, and efficient prosthetic control interface that can support versatile ADLs beyond essential hand functions, this paper is based on a high-quality HD-sEMG database that includes 65 isometric hand gestures with different degrees of freedom (DoFs) recently published in the scientific data of Nature [35]. The movements consist of 16 1-DoF finger and wrist gestures, 41 2-DoF compound gestures of fingers and wrist, and eight multi-DoF gestures of grasping, pointing, and pinching. The database was collected from 20 healthy participants, 14 males and 6 females, with wide-ranging ages between 25 and 57 years old (mean: 35 years old). We only use the signals from 19 subjects because the data from subject 5 is not available. The HD-sEMG signals were recorded using a Quattrocento (OT Bioelettronica) biomedical amplifier system through two 8×8 electrode grids (a total of 128 channels) with a 10mm inter-electrode distance, at a sampling rate of 2048 Hz. The two grids were positioned on the dorsal (outer forearm) and the volar (inner forearm) of the upper forearm. The recording was performed in a differential manner, where the channel *i* signal is the signal difference between electrode *i*+1 and electrode *i*, to reduce common-mode noise. Each subject was asked to perform each gesture for five repetitions before switching to the next one. Each repetition lasts for five seconds, followed by an equal-duration rest. Fig. 1 shows muscle-activity heatmaps from the two 8×8 electrode grids (inner and outer forearm) for the best-performing subject. Due to space limitations, we only show 16 out of the 65 gestures, and choose the simplest most visually intuitive examples. We show the heatmaps for the two grids, for a total of 32 heatmaps. It can be observed that for Movement 2 “ring finger: bend”, which is an extension of the little finger, more muscle activity is observed on the outer forearm (which contains the extensors) than on the inner forearm. Independent forces from each finger and the wrist were utilized to assist the temporal relabeling in aligning the movement labels with the segments of the hand gestures once they have reached a plateau. This paper uses the labels before the temporal adjustment to include the transient phase.

### B. Data Preprocessing

In this work, we define the length of the transient phase by averaging the corresponding force signals of each gesture across all subjects. The HD-sEMG signals of each repetition have been truncated after 0.5 secs to capture the computed transient phase average. Fig. 2 shows the 0.5-second transient phases (indicated by dashed lines) of the corresponding force signals of Movement 1 (1-DoF), Movement 26 (2-DoF), and Movement 61 (multi-DoF). We then scale up the signal magnitudes using Min-Max normalization only based on training data, followed by Mu-law transformation [36] on each data scalar in a logarithmic and nonlinear manner. Mu-law transformation is applied as can be seen in (1) to enhance the discriminability of the information among channels.

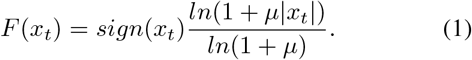

In (1), *x_t_* denotes each data scalar and *μ* = 2048. We conduct signal windowing and evaluate the effect of varying window sizes, following the real-time implementation standards in myoelectric control [34],[37]–[40]. We investigate sliding window sizes of 100ms, 200ms, and 300ms with the same step size of 10ms. Each short window is a data point for training the model. As a result, the model input has a shape of (sampling rate*window size) ×128. 128 is the number of channels (two 8×8 grids). Thus, for a 200ms window size, only 20 minutes of calibration/training data is fed to the model for each subject. It is commendable that a 65-class model can work with such little data, enhancing the practicality and reducing the need for extensive calibration.

## III. MODEL STRUCTURE

Based on our previous research and recent literature, it should be mentioned that for a large number of gestures and the steady-phase of contraction, deep neural networks can achieve high performance when given large datasets. However, deep structures and the need for large datasets are two primary factors leading to complex model architectures and long training times. Motivated by this issue, in this paper we propose heterogeneous temporal dilation, for the first time, aiming at adding longer, nonlinear, and more diverse temporal reach to the LSTM model. This paper also proposes one new degree of freedom, dilation focus, to the model structure, indicating the skewness of the connection density of the dilated LSTM cells on each layer.

### A. Regular Baseline Model and Dilated Baseline Model

We compare the model performance of the proposed heterogeneously dilated model with two baseline models: a regular LSTM model (see Fig. 3a) and a homogeneously dilated LSTM model. (See Fig. 3b). The study using the regular baseline model evaluates the effect of any dilation, while the study on the dilated baseline model compares the effect of different temporal dilation strategies (homogeneous vs. heterogeneous). For consistency, the regular baseline model consists of four LSTM layers, each having the number of LSTM cells equal to window size×sampling rate (e.g., 409 LSTM cells for a 200ms window) and 128 hidden units. The 128 hidden units of the last LSTM cell of the fourth LSTM layer are fed into the classifier (i.e., a fully connected neural net which fuses the decoded information for gesture prediction). The classifier contains three fully connected layers, sequentially including 64, 32, and 65 nodes, to conduct gesture prediction. The dilated baseline model has a similar architecture to the regular baseline model, but the 3rd-order homogeneous dilation is injected into the LSTM layers. Refer to our previous work [26] for more details on the homogeneous model and the aggressiveness of temporal dilation. Early stopping (a common technique to prevent overfitting in the literature [20],[41],[42]) with a patience factor of 30 is used. This means that the model will stop training after 30 iterations past the point at which the accuracy has plateaued.

### B. Heterogeneous Dilation and Dilation Focus

Compared with the homogeneous dilation that has vertical aggressiveness within each layer, in heterogeneous dilation, we examine different aggressiveness horizontally within the second layer. The number of skipped LSTM cells between two connected cells exponentially increases/decreases, determined by the dilation focus. In a left-focused model (see Fig. 4a), the model is divided into three equal-length time segments. The number of skipped cells (denoted as *N_k_*) of each time segment can be derived from an exponential function shown in (2).

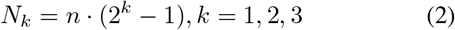

*k* denotes the k-th time segment, and *n* represents the maximum number of skip connections given the time segment. In a right-focused model (see Fig. 4c), the model is also segmented into three equal parts on the time axis. The number of the skipped cells of each time segment can be calculated from the same exponential function but with k in the reverse order. In a middle-focused model (see Fig. 4b), we first find the median cell of each layer, and then divide the LSTM model into two submodels, each having three equal-length time segments (one-sixth of the window size). The submodel on the left is equivalent to a right-focused dilated model, whereas the submodel on the right is equivalent to a left-focused dilated model. Early stopping with a patience factor of 30 is again used.

## IV. Experiments and Results

### A. Experiment Models

Following the previously explained model structures, we perform a comprehensive analysis on five LSTM-based models listed in Table I. Model 1 is a regular 4-layer LSTM network. Model 2 adds 3rd-order homogeneous dilation, skipping 7 out of every 8 cells on the second layer. (Refer to [26] for details on the upper layers.) Based on model 2, we extend to models 3-5, where we replace the homogeneously dilated second layer with the three versions of heterogeneous dilation. We adapt the heterogeneous dilation only on one layer because experiments showed that applying dilation of the same focus on too many layers results in an overall condensing of information in one area and too much loss in the other areas, therefore affecting the performance. A left-focused dilation is used on the second layer of model 3, a middle-focused dilation is used on model 4 and a right-focused dilation is used on model 5. We train user-specific models for each of the 19 subjects.

**TABLE I:** Model descriptions.

| ID | Name |
| --- | --- |
| 1 | 4-Layer, Regular Baseline Model |
| 2 | 4-Layer, Dilated Baseline Model |
| 3 | 4-Layer, Heterogeneous Dilation, Left Focus |
| 4 | 4-Layer, Heterogeneous Dilation, Middle Focus |
| 5 | 4-Layer, Heterogeneous Dilation, right Focus |

To evaluate the model generalization and performance legitimacy on new data, we conduct k-fold (k=5) crossvalidation. We hold out one repetition for testing and use the other four for training. The average results of the cross-validation are reported in Tables II and III and used for the box plots in Fig. 5.

**Fig. 5:**
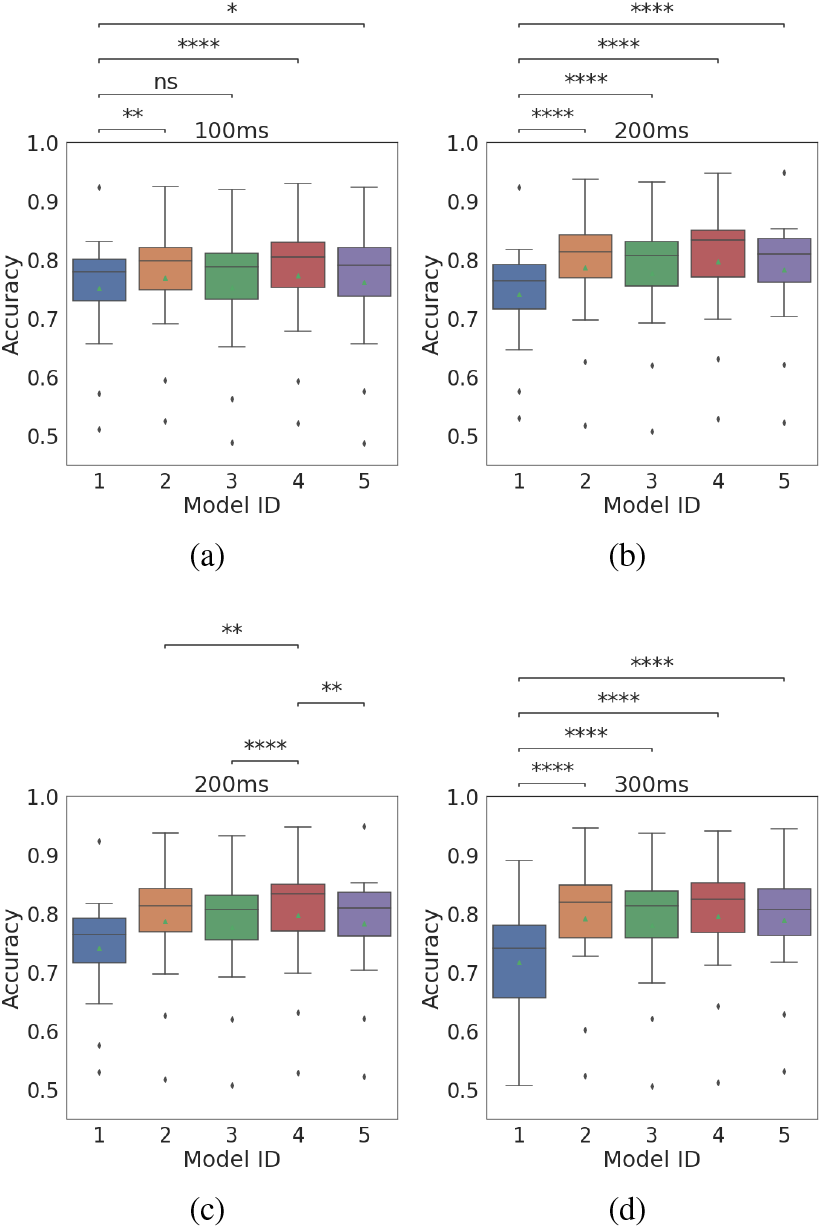
(a) Accuracy box plots of models using 100ms window size, (b) 200ms window, (c) 200ms window comparing the dilated versions (d) 300ms window.

**TABLE II:** Model accuracy.

| Model \ Window | 1 | 2 | 3 | 4 | 5 |
| --- | --- | --- | --- | --- | --- |
| 100ms | 77.9% | 79.8% | 78.8% | **80.4%** | 79.0% |
| 200ms | 76.4% | 81.3% | 80.7% | **83.3%** | 81.0% |
| 300ms | 74.1% | 82.0% | 81.3% | **82.4%** | 80.7% |

**TABLE III:**
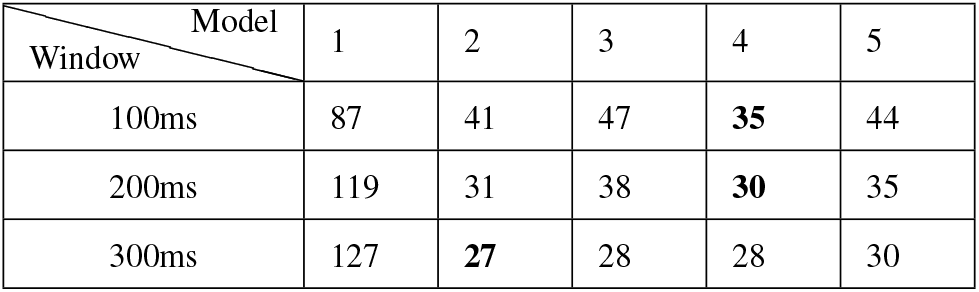
Number of converge iterations.

| Model \ Window | 1 | 2 | 3 | 4 | 5 |
| --- | --- | --- | --- | --- | --- |
| 100ms | 87 | 41 | 47 | **35** | 44 |
| 200ms | 119 | 31 | 38 | **30** | 35 |
| 300ms | 127 | **27** | 28 | 28 | 30 |

### B. Results and Statistical Analysis

We perform statistical analysis on all previously mentioned models across all 19 subjects. We perform D’Agostino-Pearson test for normality, which compares the results using paired t-tests. The significance threshold for the p-value is 0.05. We apply Bonferroni correction to the observed p-values to reduce the probability of false positives. Markers are used to denote corrected p-value ranges as following: (a) The ns marker (for not significant) denotes 0.05 to 1; (b) * denotes 0.01 to 0.05; (c) ** denotes 0.001 to 0.01; (d) *** denotes 0.0001 to 0.001; and (e) **** denotes smaller than 0.0001. Fig. 5 shows box plots of the different model prediction accuracies with the comparison markers. As can be seen, the dilated models are consistently performing better than the base models, demonstrating the power in performance of the dilation models. Among the dilated versions, the middle-focused heterogeneous dilation model (model 4) shows the best result. Table II lists the median accuracy of each model for different window sizes. We achieve the best accuracy of 83.332%, using the middle-focused structure and the 200ms window size. Fig. 5c shows the t-test results between the best-performing (middle-focused) model and other dilated models. It can be observed that the middle-focused model has a statistically significantly higher performance. This evidence shows that our proposed structure has the potential to increase accuracy while enhancing the model’s generalizability and adaptability on transient-phase data.

Similarly, we compare the number of iterations for convergence in training the model, which we define as the number of iterations required for the validation accuracy to reach 95% of the final best accuracy using the same method previously presented. We show in Table III that the dilated models require less iterations to converge. In particular, for models with window sizes of 100ms and 200ms, the middle-focused model achieves fastest convergence.

Fig. 6 compares training validation accuracies between the base model and the middle-focused heterogeneous dilation model with a 200ms sliding window for all subjects. The plot shows the progression of accuracy with training iterations. We can see that the proposed heterogeneous dilation model brings significant and consistent improvements in accuracy, convergence speed, and smoother convergence patterns.

**Fig. 6:**
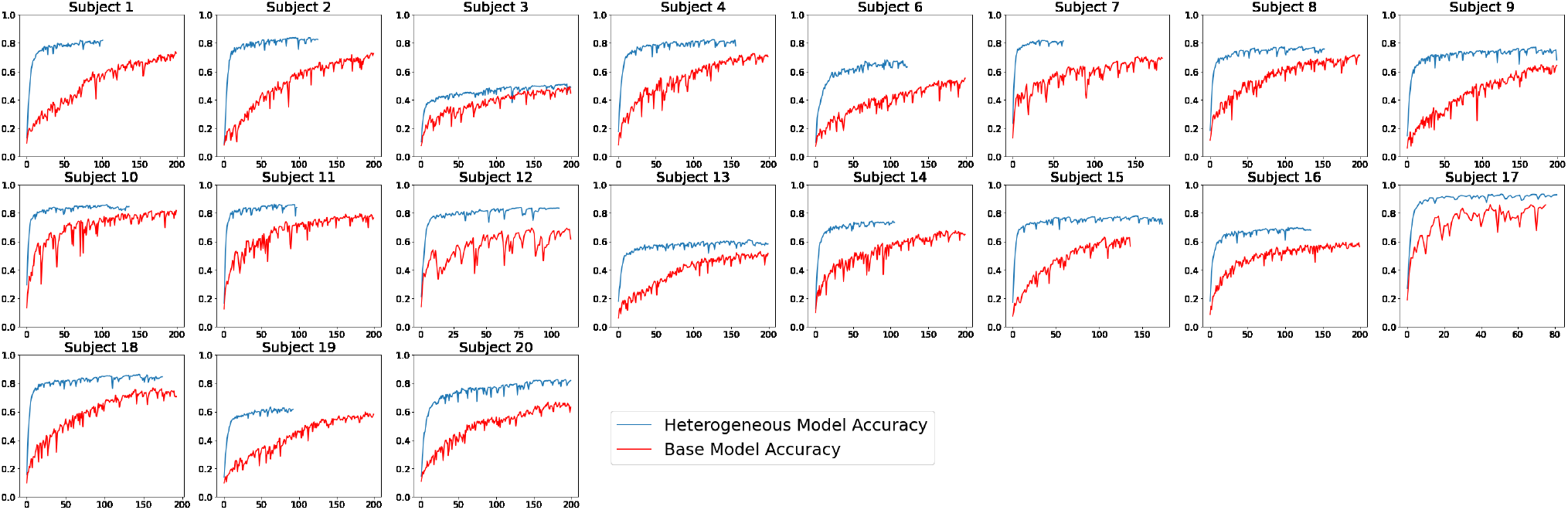
Validation accuracy per iteration (blue line is the proposed and red line is the conventional technique). (Subject 5 missing from the online database.)

## V. Comparative Study

Here we conduct a comparative study to compare the proposed heterogeneously dilated model with conventional non-sequential Deepnets that have been most commonly used in the literature [10]–[16],[43],[44]. This comparative study emphasizes the importance of sequential modeling in capturing the underlying temporal dynamics. In addition, a comparison of our proposed temporally dilated model and common sequential modeling technique, i.e., LSTM, was presented in Sect. IV (Experiments and Results) to highlight the benefits of our proposed architecture.

Thus, in this section, we compare our best (middle focused) model with a CNN and a Multilayer Perceptron (MLP). All models use a window size of 200ms. The CNN model consists of two CNN blocks, each having a convolutional layer, a batch normalization layer, and a Parametric Rectified Linear Unit activation function. The first convolutional layer has 16 filters and the second has 24 filters. Each layer has a kernel size of 15×5. A maxpooling layer with a kernel size of 2×2 is defined between CNN blocks. The outputs of the last CNN block are flattened and fed to a two-layer fully connected classifier for gesture prediction. The tested MLP model has 128 nodes on the hidden layer. The results are shown in Table. IV.

**TABLE IV:**
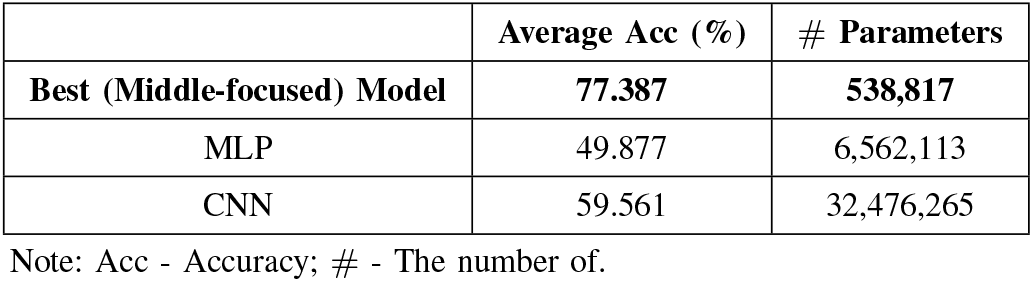
Results for comparing the proposed heterogeneously dilated LSTM model with conventional Deepnets.

|  | Average Acc (%) | # Parameters |
| --- | --- | --- |
| **Best (Middle-focused) Model** | **77.387** | **538,817** |
| MLP | 49.877 | 6,562,113 |
| CNN | 59.561 | 32,476,265 |

Note: Acc - Accuracy; # - The number of.

### Observation 1

With the limited window size and limited data from the transient phase, both CNN (with 59.561% accuracy) and MLP (with 49.877% accuracy) fail in the gesture prediction task.

### Observation 2

The proposed model has a level of information modeling and compactness that is not possible to achieve by CNN or MLP. It should be added that the trainable parameters of CNN are >60 times more than the proposed model, and the number of trainable parameters of the MLP are >12 times more than the proposed model.

## VI. Conclusion

This paper proposes a nonlinear temporal dilation, named “heterogeneous dilation”, into the LSTM layers. We have shown that the proposed structure significantly improves the training times and convergence speeds (>20 times faster) and boosts the accuracy when predicting 65 diverse gestures, compared with a non-dilated counterpart LSTM. This paper brings research one step closer to real-time implementation of prosthesis control by training the proposed model only on the transient phases, using just 10% of information at the beginning of each repetition. Moreover, the conducted study on the impact of varying window sizes has found that our proposed model achieves state-of-the-art performance when using a sliding window size of 200ms, which is shorter than the real-time implementation requirement of 300ms. The introduction of dilation focus to the proposed model adds another novel degree of freedom into the structure, shifting the model focus to prioritize the deep observations and hidden states of a particular segment of information. Hence, the heterogeneously dilated model becomes more robust, agile, and adaptable to various tasks. The fast convergence of the proposed model opens the door for ubiquitous outside-the-lab applications and for researchers who do not have access to high-performance computers.

In this work, we evaluate our heterogeneously dilated model on a large variety of HD-sEMG signals that can capture the varying neurophysiological features of 19 able-bodied subjects of different demographics and biomechanics, demonstrating that our model is adaptable and robust. The authors would like to highlight and confirm that even though we utilize an inclusive dataset to capture variability in human neurophysiology, the neurophysiology of the healthy population does not reflect the amputees’, which vary based on the nature of surgeries and underlying causes. Hence, as part of our future work, we will collect data from amputees to translate the performance of our model to practical applications. Also, this paper applies heterogeneous dilation and dilation foci in user-specific hand gesture prediction, following the convention in the literature. Leveraging our work in generalization and temporal dilation, proposing a heterogeneously dilated and generalized model for hand gesture prediction in upper-limb prosthetic control is another future line of our research. More details about our preliminary work on generalization can be found in [25].

